# Ca^2+^-driven cytoplasmic backflow secures spindle position in fertilized mouse eggs

**DOI:** 10.1101/2024.02.05.578955

**Authors:** Takaya Totsuka, Miho Ohsugi

## Abstract

Fertilization triggers hours-long Ca^2+^ oscillations in mammalian eggs, but the effects of repeated Ca^2+^ surges remain unclear. Here, we investigate spindle dynamics and its relationship with cytoplasmic streaming in fertilized mouse eggs. The spindle, initially parallel to the plasma membrane, rotates vertically, in accordance with previously reported results using artificially activated eggs. Intriguingly, it transiently reverses its rotation direction in synchrony with Ca^2+^ oscillations, regardless of artificially altered frequency. This effect results from cytoplasmic streaming, initially moving from spindle to egg center, displaying a Ca^2+^-dependent backflow. Streaming also impacts spindle positioning, balancing spindle rotation and cortical localization maintenance. We provide evidence that Ca^2+^-dependent cortical myosin II activation causes actomyosin contraction, leading to transient streaming towards non-contracting actin cap regions overlaying chromosomes. Our findings underscore the role of Ca^2+^ oscillations in maintaining spindle position in fertilized eggs, thereby ensuring highly asymmetric division and preservation of maternal stores in zygotes.

## Main

In vertebrates, eggs are arrested at metaphase II (Meta-II) while awaiting fertilization. The fusion between sperm and egg induces an elevation in free cytoplasmic Ca^2+^, triggering release from Meta-II arrest and the onset of Anaphase II (Ana-II)^1–4^. Consequently, one set of segregated chromosomes is extruded into a smaller cell, known as the second polar body (PB2). The other set of chromosomes in the larger cell contribute to development together with the sperm-derived chromosomes^5^. This process is conserved among vertebrates. However, mammals exhibit unique features, such as the delay in pronuclear (PN) formation that takes over an hour following chromosome segregation^6–10^. Furthermore, in mouse Ana-II eggs, the spindle, initially parallel to the plasma membrane, must rotate perpendicularly^5,11,12^. In parthenogenetically activated eggs, the inward cytoplasmic streaming, generated in an actomyosin dependent manner, aids in rotating the Ana-II spindle. However, it remains unknown how the Ana-II spindle rotates in fertilized eggs while retaining its cortical localization, resisting the inward streaming.

Most non-mammalian species present a single or few waves of Ca^2+^ transients after fertilization, while mammals exhibit repeated Ca^2+^ oscillations over several hours until pronuclear formation^13^. However, the initial Ca^2+^ transient triggers primarily egg activation events, including meiotic resumption and polyspermy blocking^14–18^ and the biological importance of continued Ca^2+^ oscillations during the long Ana-II remains unclear.

In this article, we address these issues by developing a live-imaging method for mouse in vitro fertilized (IVF) eggs. Our findings indicate that each Ca^2+^ oscillation induces cortical actomyosin contraction, generating transient cytoplasmic backflow. This repetitive inversion of cytoplasmic streaming helps maintain the subcortical localization of the rotating Ana-II spindle, propelling it back to the cortex when displaced. Our results highlight the importance of mammalian-specific Ca^2+^ oscillations in retaining spindle position to form a small polar body and a large egg, preserving maternal stores.

## Results

### Rotating Ana-II spindle in IVF eggs shows periodic, transient decrease in rotation angle

To investigate spindle rotational movement within fertilized mouse eggs, we first established a method for live imaging of the spindle dynamics throughout the process of IVF. We prepared a glass-bottom dish with two nearby medium drops, one with a Meta-II egg expressing EGFP-tagged α-tubulin and mRFP1-tagged histone H2B, and the other with sperm, all covered with liquid paraffin. Before placing the eggs in the medium drop, an opening in the zona pellucida for sperm entry was made. The egg was positioned in the medium drop so that the long axis of the Meta-II spindle aligned parallel to the focal plane at the equatorial plane of the egg, and then stabilized using a holding pipette. After initiating live observation, two drops of medium were connected for insemination (Fig. 1a). Confocal images, covering the entire short axis of the Meta-II spindle, were collected every 1 μm along the z-axis for a total depth of 15 μm. Using this imaging protocol, we captured the dynamics of the chromosomes and spindles from Meta-II to the extrusion of PB2 continuously (Fig. 1b and supplementary Video 1). This indicates that chromosome segregation and spindle rotation occurred in the xy-plane, where the long axis of the Meta-II spindle lies, with minor movement along the z-axis. This allowed us to analyze spindle dynamics in detail using z-stack projection images.

**Fig. 1.**
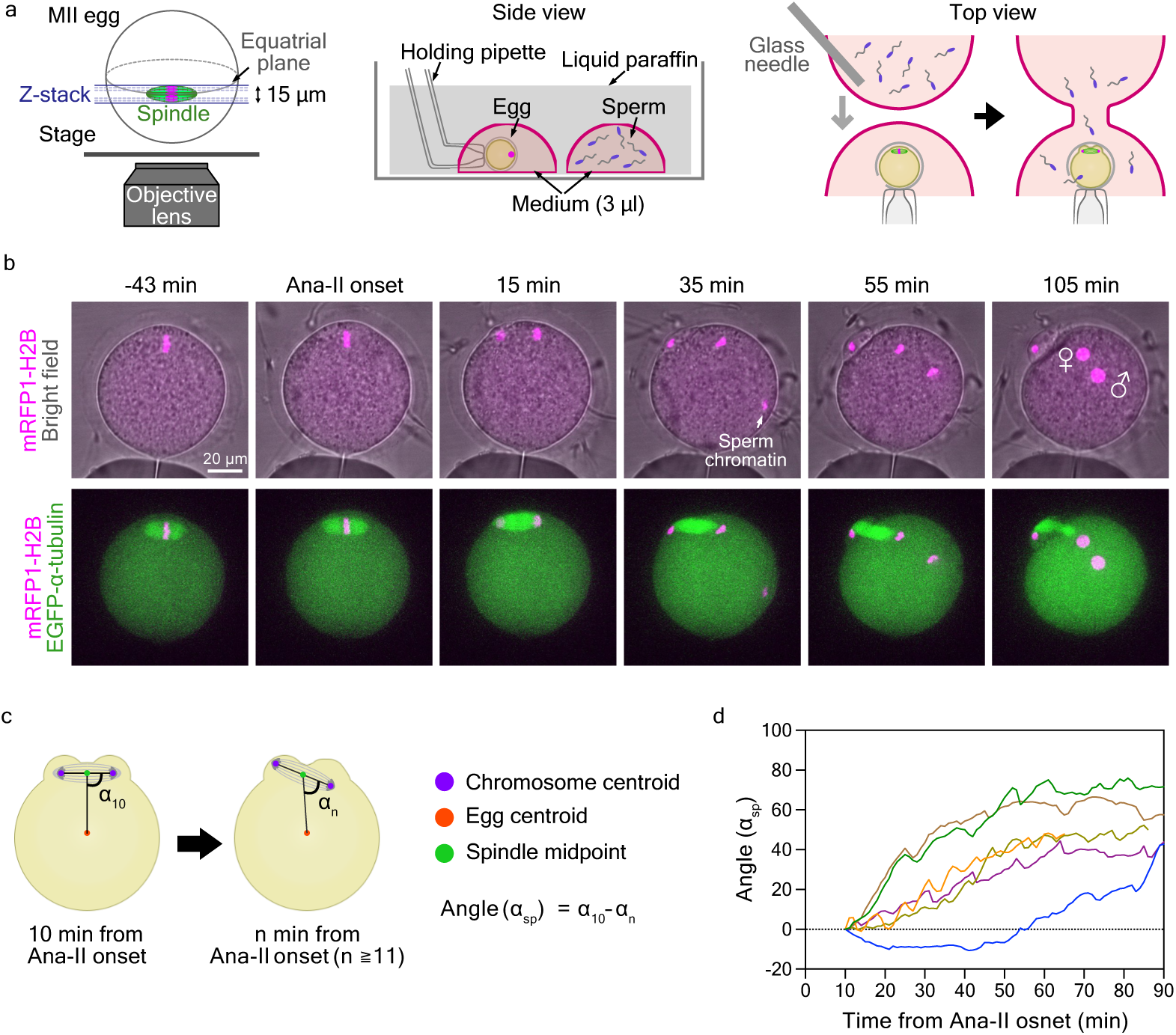
Spindle dynamics in IVF eggs. **a,** Schematics showing live imaging method for IVF eggs. Confocal images of z-slices covering the entire width of the spindle short axis in orientation-adjusted Meta-II eggs (left). Two medium drops, one containing a Meta-II egg held by a pipette and the other with sperm, were prepared on a paraffin-covered glass-bottom dish (middle). Two drops of medium were connected for insemination (right panel). **b,** Representative time-lapse images showing chromosomal and spindle dynamics in IVF eggs expressing mRFP1-tagged histone H2B (magenta) and EGFP-tagged-α-tubulin (green). The numbers above the top panels indicate the time before and after the onset of Ana-II. **c,** Schematics showing the quantification of the spindle rotation angle. **d,** Change over time in spindle rotation angle for six IVF eggs.

The rotational movement of the Ana-II spindle was analyzed using time-lapse images taken every minute. In each image, the midpoint of the centroids of the separated chromosomes was defined as the spindle midpoint; and the angle formed by the line connecting the chromosome centroids and the line connecting the spindle midpoint and the centroid of the egg cell was defined as the angle (α) (Fig. 1c). We set the angle (α) in the image taken 10 min after the onset of Ana-II (α_10_) as the standard and plotted subsequent changes in angle as the spindle rotation angle (α_sp_). Overall, the spindle angle increased over time, aligning with previously reported results using parthenogenetically activated eggs^19,20^. However, most IVF eggs exhibited a periodic, transient decrease in the rotation angle (Fig. 1d).

### Ca^2+^ oscillations cause repeated transient inversions in the direction of spindle rotation

A decrease in the rotation angle was observed every 5–15 min, although the interval varied among eggs. This pattern is similar to the interval between transient increases in Ca^2+^ in fertilized mouse eggs^21^. As expected, simultaneous observation of spindle dynamics and intracellular Ca^2+^ changes in IVF eggs revealed that when the Ca^2+^ concentration was low, spindles rotated in a positive direction; however, immediately after a Ca^2+^ transient, the direction of spindle rotation changed rapidly and temporarily switched to a negative direction (Fig. 2a,b and expanded Fig. 1a and Supplementary Video 2, top panels). A correlation between the temporal reversal of the rotational direction of the spindle and the Ca^2+^ transient was also observed in eggs activated by Sr^2+^ (hereafter referred to as Sr^2+^-activated eggs), which exhibit Ca^2+^ oscillations similar to fertilized eggs^22^ (Fig. 2c,d and Expanded Data Fig. 1b and Supplementary Video 2, bottom panels). These results demonstrate that the periodicity of spindle rotational movement coincides with that of Ca^2+^ oscillations independent of sperm-derived factors.

**Fig. 2.**
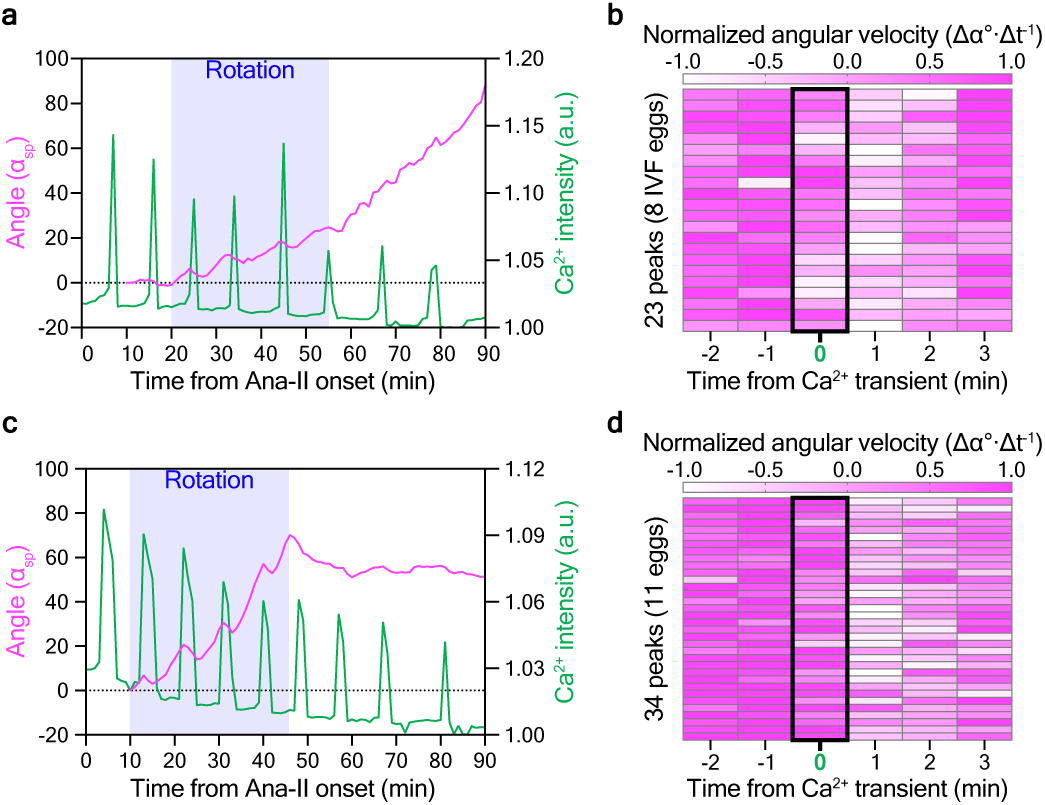
Direction of the spindle rotation transiently reverses coincidence with the Ca^2+^ transient. **a,c,** Graphs showing the spindle rotation angle (magenta) and relative intensity of cytoplasmic Ca^2+^ levels (green) in representative IVF **a**, or Sr^2+^-activated **c** eggs. **b,d,** Heat maps showing the normalized angular velocity (Δt = 1 min) of spindle rotation for several minutes before and after the Ca^2+^ transient in IVF eggs (23 peaks, 8 IVF eggs) **b**, or in Sr^2+^-activated eggs (34 peaks, 11 eggs) **d**. The timing of the Ca^2+^ transient is enclosed by a black square.

Next, to examine the impact of changes in Ca^2+^ oscillations on the spindle rotation, we first tried to increase the frequency of Ca^2+^ oscillations using human-PLCζ^23–25^. Exogenously expressed human-PLCζ increased the frequency of Ca^2+^ oscillations in a dose-dependent manner, but only either before or after spindle rotation. We then combined human-PLCζ expression with Sr^2+^ activation and succeeded in triggering high-frequency Ca^2+^ oscillations during spindle rotation (Fig. 3a,b). A correlation between the temporal change in the rotation direction and the Ca^2+^ transient was observed even in eggs with increased frequency of Ca^2+^ oscillations (Fig 3b,c and Extended Data Fig. 1c). Second, thapsigargin, an inhibitor of sacro-ER Ca^2+^-ATPase (SERCA)^26,27^, was added to the medium during the early Ana-II to inhibit Ca^2+^ oscillations (Fig. 3d). Following this inhibition, the spindle exhibited smooth rotational movement, without any transient reversal in its rotation direction (Fig. 3e and Extended Data Fig. 1d). Together, these results indicate that elevated cytoplasmic Ca^2+^ levels reverse spindle rotation.

**Fig. 3.**
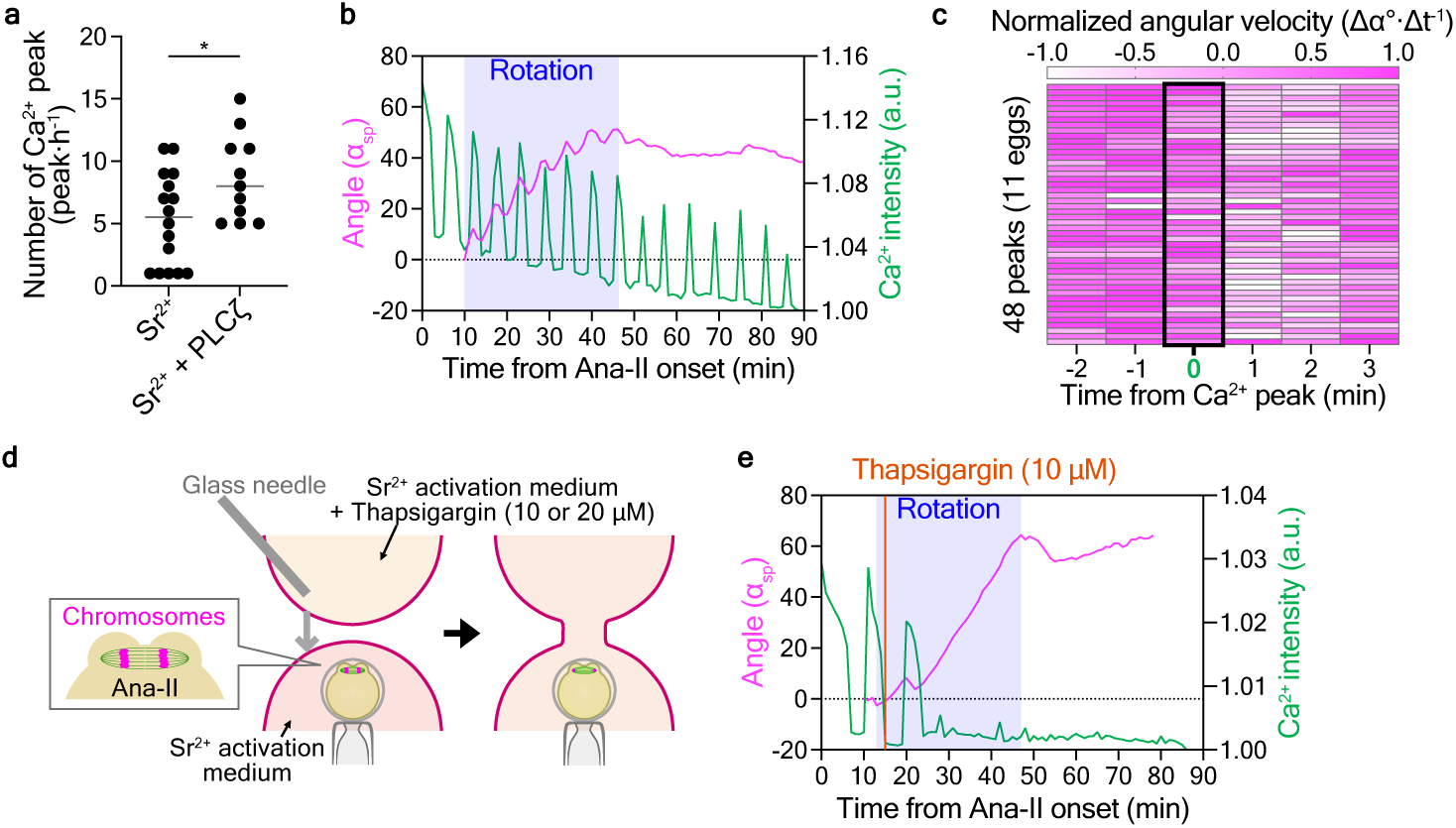
Inversions in the direction of spindle rotation depends on Ca^2+^ transients. **a,** The frequency of Ca^2+^ oscillations during 10 to 70 min after Ana-II onset in Sr^2+^-activated eggs expressing or not human-PLCζ. The gray bars indicate average values. *, p < 0.05; Welch’s t test. **b,** A graph showing the spindle rotation angle (magenta) and relative intensity of cytoplasmic Ca^2+^ levels (green) in representative Sr^2+^-activated eggs expressing human-PLCζ. **c,** Heat map showing the normalized angular velocity (Δt = 1 min) of spindle rotation for several minutes before and after the Ca^2+^ transient in human-PLCζ expressing Sr^2+^-activated eggs (48 peaks, 11 eggs). The timing of the Ca^2+^ transient is enclosed by a black square. **d,** Schematic of thapsigargin treatment for Ana-II eggs during live observation. **e,** Graph showing the spindle rotation angle (magenta) and relative intensity of cytoplasmic Ca^2+^ levels (green) in a representative thapsigargin-treated egg. Orange bars indicate the time of thapsigargin addition.

### Ca^2+^-induced outward cytoplasmic streaming causes a reversal of spindle rotation

To further investigate the Ca^2+^-dependent changes in the direction of spindle rotation, we analyzed cytoplasmic streaming, considering its supposed role in driving spindle rotation during Ana-II^19,20^. Using particle image velocimetry (PIV; see Materials and Methods), we revealed that in Ana-II, inward cytoplasmic streaming was observed as previously reported^19^, but immediately after the Ca^2+^ levels peaked, the orientation of the streaming reversed transiently to outward (Fig. 4a-c and Expanded Data Fig. 2a and Supplementary Video 3). The directions of spindle rotation and cytoplasmic streaming always coincided: when cytoplasmic streaming was inward or outward, the spindle rotated in the positive or negative direction, respectively (Expanded Data Fig. 2b). To clarify the causal relationship between directional changes of cytoplasmic streaming and spindle rotation, we utilized a method of partial spindle disruption using low-dose nocodazole. This approach halted chromosome segregation and spindle rotation without compromising meiotic resumption upon egg activation^28^. The low-dose nocodazole did not affect Ca^2+^ oscillations, and the cytoplasmic streaming was oriented outward immediately after each Ca^2+^ transient, even in the absence of spindle rotation (Fig. 4d-f and Expanded Data Fig. 2c and Supplementary Video 4). These results demonstrate that elevated cytoplasmic levels of Ca^2+^ induce the inversion of cytoplasmic streaming, thereby changing the direction of spindle rotation.

**Fig. 4.**
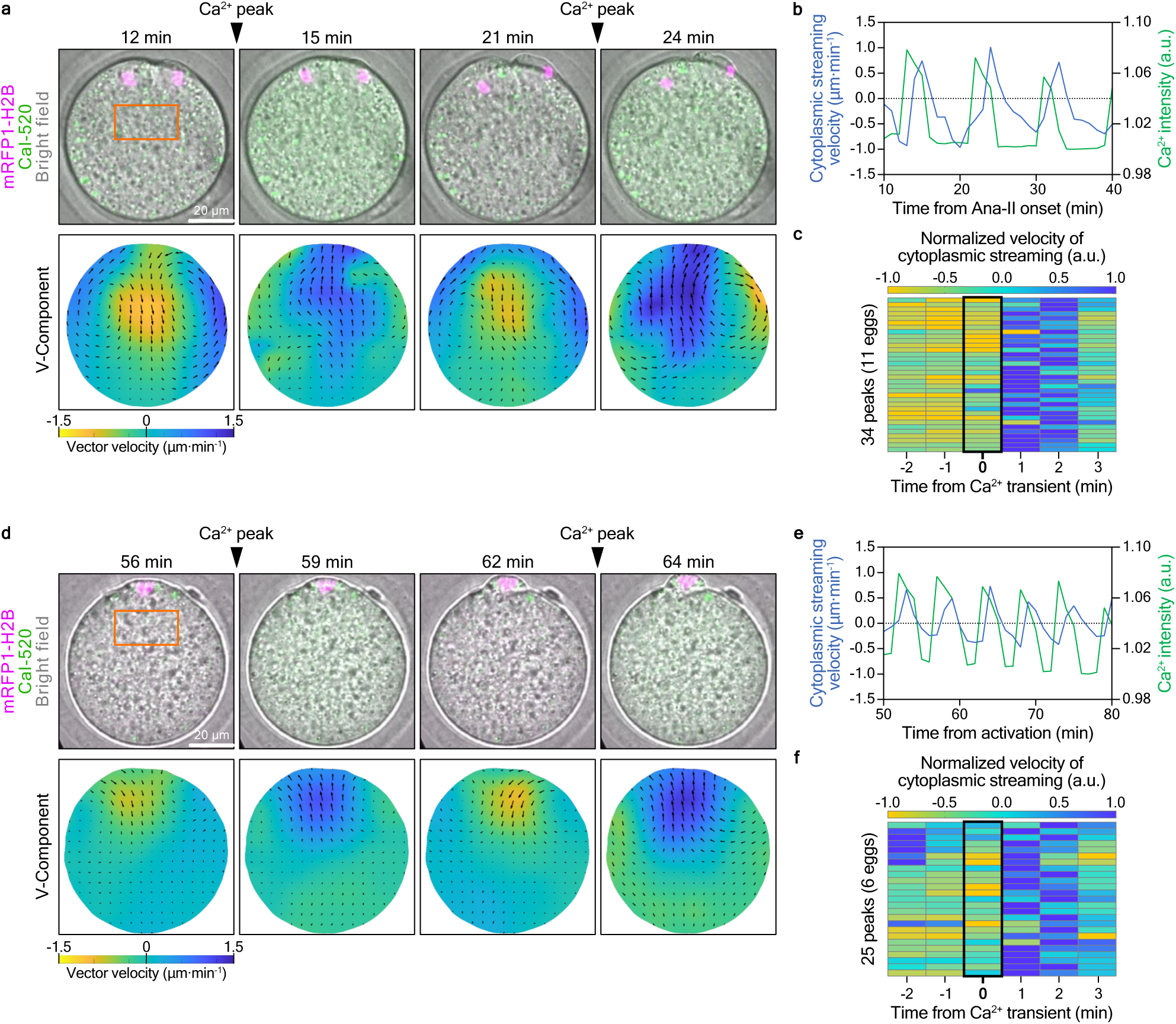
Cytoplasmic streaming inverts immediately after Ca^2+^ levels peak. **a,d,** Representative images showing chromosomes (magenta) and cytoplasmic Ca^2+^ levels (green) in a Sr^2+^-activated egg **a**, or a Sr^2+-^-activated egg treated with 0.08 µg/mL nocodazole **d** at a time point before and immediately after the Ca^2+^ transient, merged with bright field (top panels). Colored heat maps of vectors indicate the direction and velocity of cytoplasmic streaming analyzed using PIV as v-component (bottom panels). The numbers above the top panels indicate the time after Ana-II onset **a**, or Sr^2+^-activation **d** (min). **b,e,** Graphs show the velocity of cytoplasmic streaming (blue) and relative intensity of cytoplasmic Ca^2+^ levels (green) in the eggs shown in **a** and **d**, respectively. The velocity of cytoplasmic streaming was analyzed within the orange frames in the upper panels of **a** and **d**. **c,f,** Heat maps showing the normalized velocity of cytoplasmic streaming for several minutes before and after the Ca^2+^ transient in Sr^2+^-activated eggs (34 peaks, 11 eggs) **c**, or in Sr^2+^-activated eggs treated with 0.08 µg/mL nocodazole (25 peaks, 6 eggs) **f**. The timing of the Ca^2+^ transient is enclosed by a black square.

### Ca^2+^ oscillations are involved in the maintenance of the subcortical localization of the Ana-II spindle and ensure the extrusion of small-sized PB2

What is the physiological importance of Ca^2+^-induced changes in cytoplasmic streaming? To address this question, we next focused on the position of the Ana-II spindle because in Meta-II-arrested eggs, both inward and outward cytoplasmic streaming affect the position of the spindle and even transports it to the egg center^29^. The position of the Ana-II spindle was analyzed by measuring the distance between the midpoint of the spindle and the centroid of the egg (Fig. 5a). In most of the Sr^2+^-activated eggs, inward cytoplasmic streaming resulted in a slight displacement of the spindle toward the interior of the egg. Conversely, when outward cytoplasmic streaming occurred, the spindle relocated toward the cortex, while consistently maintaining a certain distance from the plasma membrane (Fig. 5b,c). We also observed an egg in which the spindle unintentionally shifted away from the cell membrane during the early Ana-II stage. In this egg, the spindle gradually moved closer to the plasma membrane each time the cytoplasm flowed outward (Fig. 5d,e). These results suggest that transient, repetitive outward cytoplasmic streaming induced by Ca^2+^ oscillations is involved in maintaining the subcortical localization of the Ana-II spindle.

**Fig. 5.**
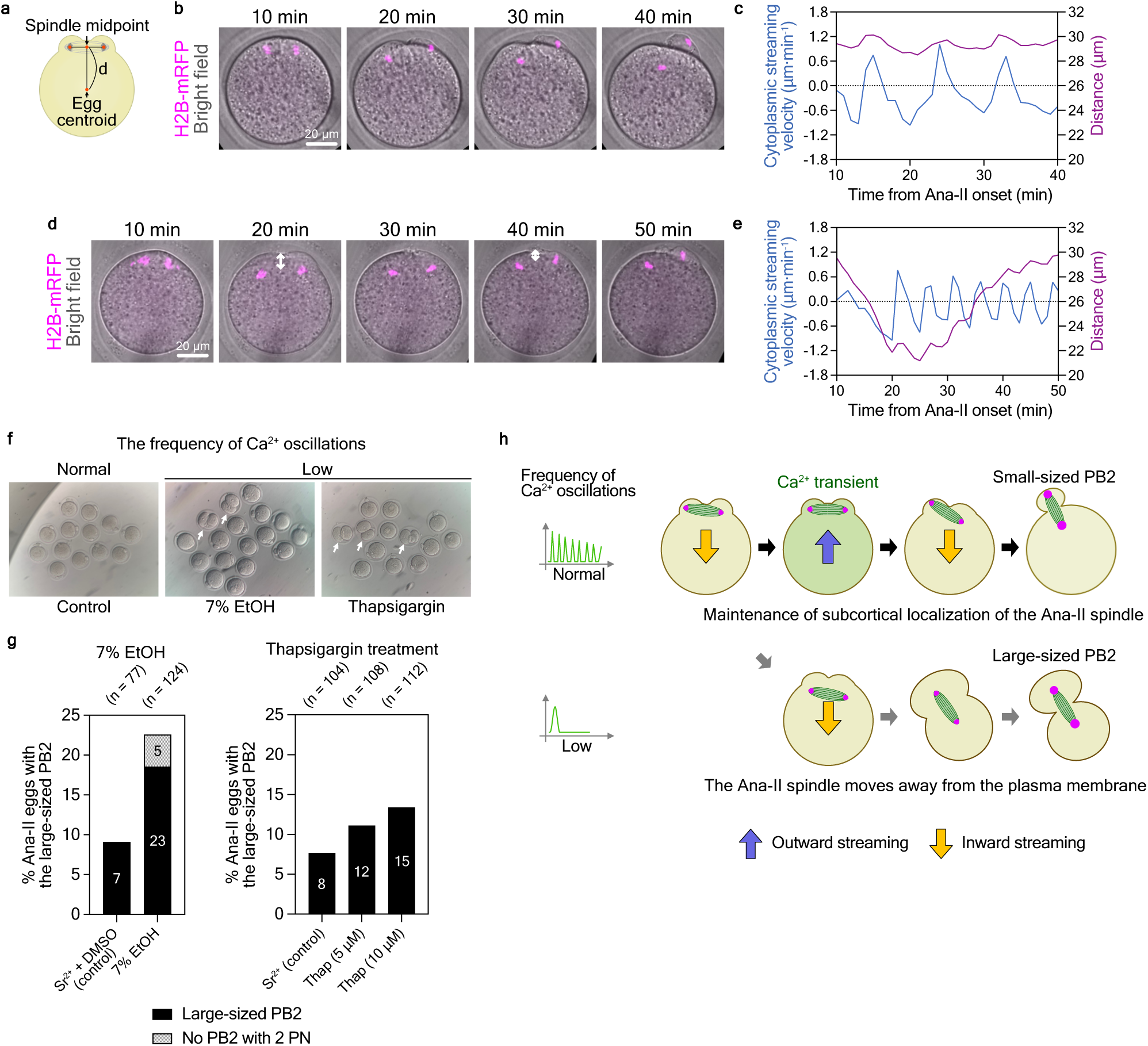
Ca2+ oscillations ensure the extrusion of a small-sized PB2. **a,** Schematic showing the quantification of the spindle position and distance (*d*) between the midpoint of the spindle and centroid of the egg. **b,** Representative time-lapse images showing the spindle position of a Sr^2+^-activated egg expressing mRFP1-tagged histone H2B (magenta). **c,** Graph showing the spindle position (purple) and mean velocity of cytoplasmic streaming (blue) for 30 min after Ana-II onset of an egg shown in **b**. **d,** Representative time-lapse images of a Sr^2+^-activated egg with spindle detachment from the plasma membrane during early Ana-II. **e,** Graph showing the spindle position (purple) and mean velocity of cytoplasmic streaming (blue) for 40 min after Ana-II onset of an egg shown in **d**. **f,** Representative images show eggs at five hours of culturing after activation with Sr^2+^ (left), 7% ethanol (middle), and Sr^2+^ followed by thaprigargin treatment (right). White arrows indicate eggs with large PB2. **g,** Ratio of activated eggs with large and no PB2 under each activation condition. The numbers in the bars indicate the number of activated eggs in each category. n: number of activated eggs observed in each experiment. **h,** Model of the mechanism that ensures subcortical localization of the Ana-II rotating spindle.

Subcortical localization of the Ana-II spindle is essential for the formation of the small PB2^30^. Therefore, to examine this hypothesis, we tested the effect of no- or lower-frequency of Ca^2+^ oscillations on the size of the PB2. First, we activated eggs with Sr^2+^ or 7% ethanol and found that 9.1% and 18.6% of the eggs, respectively, extruded a larger PB2 (Fig. 5f,g). Additionally, Sr^2+^-activated eggs were treated with thapsigargin 10 min after immersing them in the Sr^2+^- containing medium drop. This inhibited Ca^2+^ oscillations almost completely before the beginning of spindle rotation (Supplementary Video 5). In the presence of 5 or 10 µM thapsigargin 11.1% and 13.4% of eggs extruded a large-sized PB2, respectively, whereas 7.7% of the control DMSO-treated activated eggs formed a larger PB2 (Fig. 5f,g). These results suggest that Ca^2+^ oscillations change the direction of cytoplasmic streaming toward the chromosomes, thereby contributing to the maintenance of the subcortical localization of the rotating spindle (Fig. 5h).

### Cortical actomyosin contraction generates cytoplasmic flow toward the non-contracted actin cap regions

To uncover the mechanisms underlying the change in cytoplasmic streaming upon elevated cytoplasmic Ca^2+^, we first performed PIV analysis using time-lapse images of the IVF eggs, in which both maternal and paternal chromosomes appear within the same 15 µm thick z-sections. Similar to Sr^2+^-activated eggs, the direction of cytoplasmic streaming in IVF eggs was inward from the regions of the Ana-II spindle during the intervals between each Ca^2+^ peak. In contrast, immediately after Ca^2+^ levels peaked, cytoplasmic streaming was oriented toward both the maternal and paternal chromosomes (Fig. 6 and Expanded Data Fig. 3 and Supplementary Video 6). These results suggest that elevated levels of cytoplasmic Ca^2+^ change the direction of cytoplasmic streaming toward the chromosomes, rather than simply backward.

**Fig. 6.**
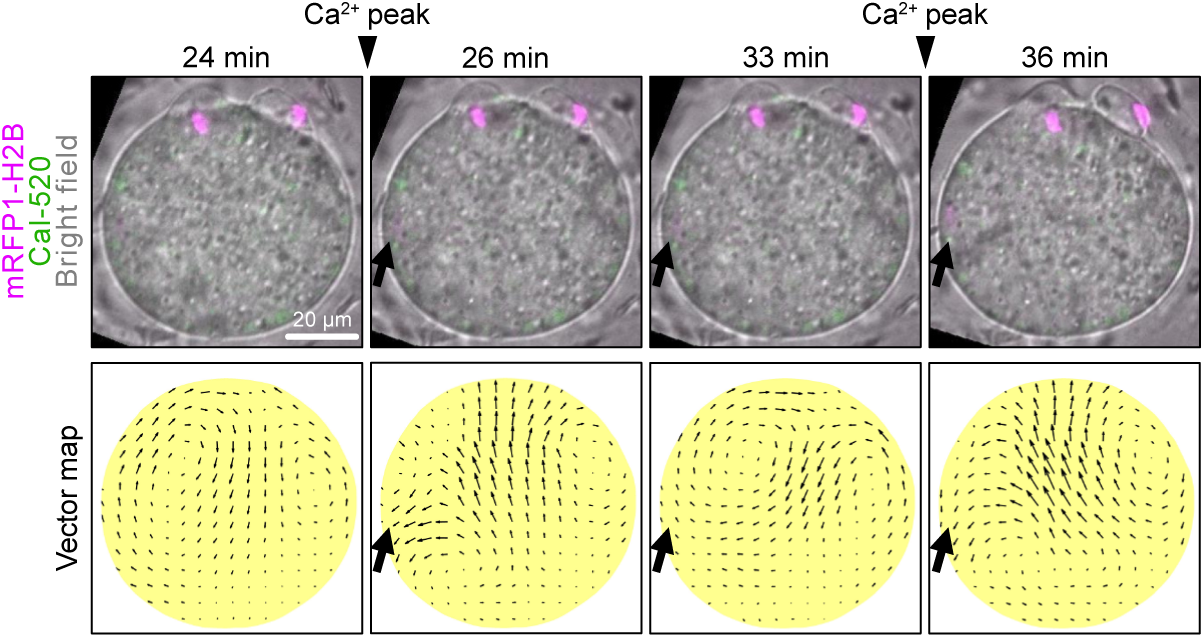
Ca2+-induced cytoplasmic streaming is oriented toward both parental chromosomes. Representative images showing chromosomes (magenta) and cytoplasmic Ca^2+^ levels (green) in an IVF egg at a time point before and immediately after the Ca^2+^ transient merged with a bright field (top panels). Vector maps indicating the direction and velocity of cytoplasmic streaming analyzed using PIV (bottom panels). Black arrows indicate the positions of sperm chromosomes. Numbers above the top panels indicate the time after the onset of Ana-II.

Given that the contraction of actomyosin has been reported to drive cytoplasmic movement in fertilized eggs^31^, we next proceeded with experiments to inhibit myosin II by adding blebbistatin 10 min after the onset of Ana-II. Ca^2+^ oscillations remained unaffected, yet the velocity of both inward and outward cytoplasmic streaming significantly decreased (Fig. 7a,b and Expanded Data Fig. 4 and Supplementary Video 7). This suggests that the activity of myosin II at disparate sites is the underlying cause of both inward and Ca^2+^-dependent outward cytoplasmic streaming. To further examine this possibility, we visualized F-actin in Sr^2+^-activated eggs and observed periodic fluctuations in F-actin intensity (Supplementary Video 8). Quantitative analysis revealed that in the cortical regions, the F-actin intensity increased concomitantly with a slight but distinct contraction of the cortical plasma membrane, outward movement of cytoplasmic streaming, and expansion of the protrusions in the actin cap regions (Fig. 7c-e). The changes in F-actin intensity within the cytoplasm contrasted and complemented the changes in the cortical regions (Fig. 7e). Previous studies demonstrate that active myosin II was localized exclusively to the regions surrounding the two protrusions overlying the segregated chromosomes and the furrow region enclosed by these two protrusions, as well as to the region surrounding the fertilization cone^19,20,31,32^. Immunofluorescence staining with an anti-phosphorylated regulatory myosin light chain (pMLC) antibody revealed that in addition to these sites, approximately 12.4% of the Sr^2+^- activated eggs exhibited cortical MLC phosphorylation (Fig. 7f and Extended Data Fig. 5a,b), whereas ethanol-activated eggs did not exhibit MLC phosphorylation at the cortex (Extended Data Fig. 5c). Moreover, in Ana-II eggs with cortical MLC phosphorylation, cortical F-actin was more pronounced than in those without cortical MLC phosphorylation (Fig. 7g), supporting the idea that cortical actomyosin contraction causes cortical actin thickening.

**Fig. 7.**
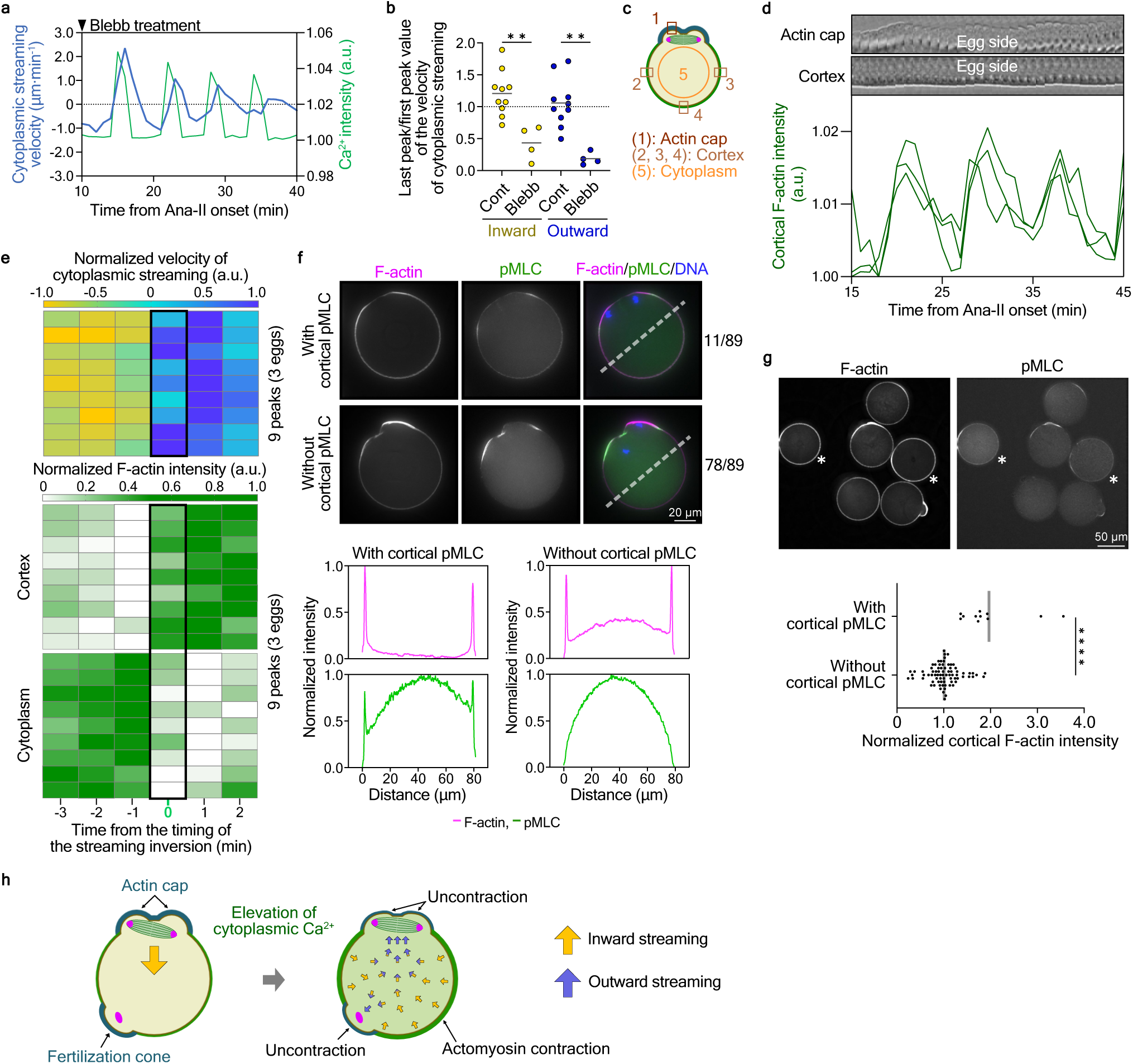
Direction of inverted cytoplasmic streaming is determined by spatial-asymmetric contractions of cortical actomyosin. **a,** Graph showing the velocity of cytoplasmic streaming (blue) and relative cytoplasmic Ca^2+^ levels (green) in a blebbistatin-treated Ana-II egg. A black arrowhead indicates the time of blebbistatin addition. **b,** Changes in inward (yellow) and outward (blue) cytoplasmic streaming at 30 min after blebbistatin treatment. The gray bars indicate average values. **, p < 0.01; Mann-Whitney test. **c,** Schematics showing the area where F-actin fluorescence intensity and plasma membrane dynamics were measured. **d,** Kymographs showing the plasma membrane dynamics of the actin cap region (region #1 in **c**) (top) and the cortical region (region #2 in **c**) (middle), and the graph showing the relative F-actin fluorescence intensity in the three cortical regions (region #2, 3, 4 in **c**) (bottom) of a representative egg. The timescales of the kymographs and graphs were consistent. **e,** Heat maps showing the normalized velocity of cytoplasmic streaming (top) and normalized F-actin fluorescence intensity in three cortical regions (region #2, 3, 4 in **c**) (middle) and the cytoplasmic region (region #5 in **c**) (bottom) in activated eggs before and after cytoplasmic streaming inversion. The minute zero indicates the timing of cytoplasmic streaming inversion (black squares). **f,** Top, representative immunofluorescent images of F-actin (magenta) and pMLC (green) in Sr^2+^-activated eggs with (top panels) and without (middle panels) cortical pMLC signals. Bottom, graphs show the fluorescence intensity profiles of F-actin and pMLC along the dashed white lines in the upper panels. **g,** Representative immunofluorescent images of F-actin and pMLC in Sr^2+^-activated Ana-II eggs. Asterisks indicate eggs with cortical pMLC (top panel). The bottom graph shows cortical F-actin signal ratios. The gray bars indicate average values. ****, p < 0.0001; Mann-Whitney test. **h,** Model of Ca^2+^-driven, chromosome-directed cytoplasmic flow.

In summary, we propose that Ca^2+^-induced contraction of cortical actomyosin pushes the adjacent cytoplasm toward the center of the cell. This merges into the streaming toward the uncontracted regions, specifically the actin cap around the maternal chromosomes and the fertilization cone around the paternal chromatin (Fig. 7h).

## Discussion

In this study, we describe a first-time detailed examination of spindle dynamics in fertilized mouse eggs. Our straightforward yet effective live imaging approach revealed two features of the Ana-II spindle rotational movement in fertilized mouse eggs. First, the spindle rotates two-dimensionally in the equatorial plane, wherein its long axis resides within the Meta-II egg. This has been assumed in previous studies using artificially activated eggs but was directly demonstrated in this study and could provide important insights into understanding the nature of forces driving spindle rotation in mouse Ana-II eggs. Second, the rotating Ana-II spindle transiently reverses its rotation direction upon cytoplasmic Ca^2+^ elevation. Repeated transient changes in the rotation angle disappeared when Ca^2+^ oscillations were suppressed by thapsigargin (Fig. 3e). This is consistent with previous reports that in ethanol-stimulated eggs, in which only a single or a few waves of Ca^2+^ transients were observed, the rotation angle of the spindle increased smoothly and with an almost constant velocity^19^.

We have uncovered, for the first time, the effects induced in eggs by mammalian-specific calcium dynamics, Ca^2+^ oscillations, triggered by sperm fusion. Our results show that throughout Ana-II, the spindle undergoes a cycle wherein it is subjected to inward cytoplasmic streaming, lasting several to tens of minutes, followed by a backflow of a few minutes that occurs in synchronization with Ca^2+^ oscillations. In Meta-II eggs, inhibition of the Arp2/3 complex induces myosin II-dependent inward cytoplasmic streaming, moving the spindle toward the egg center^29^. This implies that Ana-II spindle is constantly at risk of moving away from the cortex. Indeed, despite the centralspindlin complex anchoring the central spindle to the cortex^20^, there are occasional instances where the spindle inadvertently detaches from the cortex (Fig. 5d,e and Supplementary Video 5, bottom panels). In such cases, Ca^2+^-dependent cytoplasmic streaming toward the spindle counteracts the inward streaming and assists in maintaining cortical spindle localization. Thus, Ca^2+^ oscillations specific to mammalian eggs help mitigate the risks posed by the long-lasting Ana-II phase unique to mammals^8,10^. The Ana-II spindle rotation is essential for the proper extrusion of PB2 in mouse eggs^5,12,33^. However, in many other mammals, including humans, the long axis of the Meta-II spindle is perpendicular to the cell membrane and does not require rotation^34–36^. In contrast, regardless of the spindle orientation, the Ana-II spindle must sustain its cortical localization until the completion of cytokinesis. Therefore, we believe that the importance of Ca^2+^-dependent outward cytoplasmic streaming lies not in its ability to control spindle rotation speed, but in ensuring the subcortical localization of the Ana-II spindle and formation of a large fertilized egg^30^.

Observation of fertilized eggs revealed that the Ca^2+^-induced cytoplasmic streaming is not simply reversing the preceding flow, but is rather directed toward the membrane regions lined by the actin cap structure (Fig. 6 and Extended Data Fig. 3). Based on our results and previous observations^19,20^, we propose that inward and outward cytoplasmic streaming is driven by the continuous contraction of actomyosin near the Ana-II spindle and intermittent contractions of other cortical membrane regions due to MLC phosphorylation by myosin light chain kinase (MLCK), which is activated by Ca^2+^/CaM^14^, respectively. This model can explain the distinctive changes in Ca^2+^-dependent cytoplasmic streaming, which initially flows toward the developing PB2 as well as fertilization cone, would later transition into a fast, clear flow toward the fertilization cone after the formation of PB2^31,37^.

Cytoplasmic movement in fertilized eggs can predict subsequent developmental potential^31^. This could be attributed to the effect of Ca^2+^-independent and dependent cytoplasmic streaming on the arrangement of spindles, chromosomes, and various other organelles within the fertilized egg^38,39^. Observing the movement of cytoplasm and its impacts in zygotes of diverse mammals could further deepen our understanding of the significance of Ca^2+^ oscillations in mammals.

## Methods

### Mice

The mouse strains used in this study were purchased from Charles River Laboratories (ICR mice) and CLEA Japan (female BALB/cA and male C57BL/6J mice). ICR mice were used for egg collection. Female BALB/cA and male C57BL/6J mice were cross-bred to obtain BALB/cA × C57BL/6L F1 (CB6F1) mice, which were then used for sperm collection. All mice used in this study were 8–24-week-old. The animal experiments were approved by the Animal Experimentation Committee of the Graduate School of Arts and Sciences, The University of Tokyo (approval no. 26-29) and performed following the guidelines for animal use issued by the Committee of Animal Experiments at The University of Tokyo.

### Egg and sperm collection

Female mice (8–16-week-old) were superovulated by intraperitoneal injection of 5 IU of pregnant mare serum gonadotropin (ZENOAQ) for 48 h and 5 IU of human chronic gonadotropin (Kyoritsu Seiyaku) for 18–20 h prior to egg collection. Egg-cumulus complexes were collected in an M2 medium (M7167, Sigma-Aldrich) containing 100 µg/mL hyaluronidase to remove cumulus cells. Denuded Meta-II eggs were cultured in M16 medium (M7292, Sigma-Aldrich) covered with liquid paraffin (Specially Prepared Reagent, Nacalai tesque) at 37°C under 5% CO_2_ until mRNA injection, followed by parthenogenic activation or insemination. Spermatozoa isolated from the cauda epididymis were precultured in HTF medium (ARK Resource) covered with liquid paraffin for at least 1 h at 37°C under 5% CO_2_ before being used for insemination.

### Egg activation

For Sr^2+^ activation, we used the Sr^2+^-induced method ^40^. Meta-II eggs were placed in M16 medium supplemented with 5 mM SrCl_2_ and 5 mM EGTA and cultured at 37°C. For ethanol activation, Meta-II eggs were placed in M2 medium supplemented with 7% ethanol for 4.5 min at 25°C followed by washout and culture in M16 at 37°C. To induce high-frequency Ca^2+^ oscillations, Meta-II eggs microinjected with mRNA of human-PLCζ were inculcated for 30 min and then placed in M16 medium supplemented with 2.5 mM SrCl_2_ and 2.5 mM EGTA at 37°C.

### mRNA preparation and microinjection

mRNA was synthesized in vitro with linearized template plasmids using the RiboMax Large Scale RNA Production System-T7 (Promega) supplemented with the Ribo m^7^G Cap Analog (Promega), as described previously^41^. Template plasmids for human-PLCζ were constructed using cDNA from a pcDNA-flag-human-PLCζ-AID-EGFP-polyA. The mRNA used for microinjection were mRFP1-tagged histone H2B at 50 ng/µL, EGFP-α-tubulin at 200 ng/µL, EGFP-UtrCH at 100 to 400 ng/µL, and human-PLCζ at 10 ng/µL. A minimal amount (picoliters) of mRNA was microinjected into Meta-II eggs using a piezo-driven micromanipulator (Prime Tech) and cultured for at least 3.5 h before being subjected to live imaging.

### Live imaging

Confocal images were collected with a microscope (IX71; OLYMPUS) equipped with a spinning disk confocal system (CSU10; Yokogawa), 60x/1.30 Sil or 20x/0.85 NA oil objective lens (OLYMPUS), and a CCD camera (iXon DU897E-CSO-#BV; ANDOR) controlled by Metamorph (Universal Imaging). To observe spindle dynamics with a 60x/1.30 Sil objective lens, confocal images were collected as z-stacks at 1-µm intervals (number of optical z-stacks:16) every minute. To observe multiple eggs with a 20x/0.85 NA oil lens, confocal images were collected as z-stacks at 5-µm intervals (number of optical z-stacks: 16) every minute.

### Live imaging of IVF eggs

Removing cumulus cells causes the stiffness of the zona pellucida and prevents sperm from passing through it. Therefore, after the microinjection of mRNA, a hole was made in the zona pellucida some distance from the Meta-II spindle using a piezo-driven pipette for sperm entry through the zona pellucida, as described previously^39^. Meta-II eggs were immobilized by suctioning the position opposite the spindle with a holding pipette. Sperm cells separated using the swim-up method were placed into the HFT medium so that the sperm concentration was 2.0 × 10^6^/mL and were then cultured at 37°C under 5% CO_2_. For the live imaging of IVF eggs, we prepared two adjacent 3-µL drops of M16 medium on a Φ3.5-cm glass-bottom dish, which was covered with liquid paraffin. After adding the mRNA-microinjected eggs with the hole in the zona pellucida and a few µl of 2.0 × 10^6^/mL sperm to each medium drop, respectively, a glass-bottom dish was placed in a stage-top incubator at 37°C under 5% CO_2_. After immobilizing the orientation-adjusted Meta-II eggs using a holding pipette, insemination was performed by connecting two drops using a glass needle.

### Live imaging of parthenogenetic-activated eggs

mRNA-microinjected Meta-II eggs were washed in Sr^2+^ activation medium and transferred to 3 µL of Sr^2+^ activation medium covered with liquid paraffin on a glass-bottom dish. The orientation of the Meta-II eggs was adjusted by mouth pipetting so that the Meta-II spindle was in the equatorial plane of the eggs, with its axis parallel to the focal plane. A glass-bottom dish was then placed in the stage-top incubator at 37°C under 5% CO_2_.

### Monitoring the Ca^2+^ oscillation

To monitor the cytoplasmic Ca^2+^, Meta-II eggs were microinjected with a few picotiters of 1 mg/mL of the Ca^2+^-sensitive fluorescent dye, Cal520®-Dextran Conjugate *MW 3,000* or Cal-590^TM^-Dextran Conjugate *MW 3,000* (AAT Bioquest), along with mRNAs for the expression of the protein(s) of interest, followed by culturing for 4 hours until the start of live imaging.

### Drug treatment under live observation

Medium drops containing the drugs dissolved in DMSO were prepared as previously reported ^28^. For the thapsigargin treatment assay, two Sr^2+^ activation medium drops, one containing 10 or 20µm thapsigargin and one without thapsigargin were placed close to each other on a glass-bottom dish and covered with 4 ml liquid paraffin followed by incubation at 37°C under 5% CO_2_ for 30 to 60 min before live imaging. Eggs were placed in the thapsigargin-free medium drop and the glass-bottom dish was placed in a stage-top incubator at 37°C under 5% CO_2_. After immobilizing the orientation-adjusted eggs using a holding pipette equipped on the stage of a confocal microscope, the two drops were connected using a glass needle–5–10 min after Ana-II onset. Blebbistatin was applied 10 min after Ana-II onset to a final concentration of 100 µM, using the same method as in the thapsigargin treatment assay.

### Immunostaining

Eggs were placed in acidic Tyrode’s solution to remove their zona pellucida, followed by washout with 0.5% polyvinyl pyrrolidone/PBS and fixation in 4% paraformaldehyde for 30 min at 25°C. Fixed samples were permeabilized with 0.1% Triton-X100/PBS for 15 min at 25°C and then placed in a blocking solution (PBS containing 3% BSA and 0.1% Tween20) for 2 hours. After blocking, samples were incubated with the primary antibody, rabbit anti-phospho-myosin light chain 2 (Ser19) (pMLC) (1:100, Cell Signaling Technology #3671), for 2 hours, followed by three washes for 20 min. The samples were then incubated in the solution containing Alexa Fluor 488-conjugated goat anti-rabbit (1:500, A-11034, Invitrogen), rhodamine phalloidin (Invitrogen), and Hoechst 33342 dye (10 µg/mL, H-1399, Thermo Fisher Scientific) for 45 min, followed by three washes for 20 min. Z-stack images were collected at 1-µm intervals (number of optical z-stacks: 90) to visualize the entire eggs using a fluorescence microscope (IX70; OLYMPUS) equipped with 60x/1.30 NA Sil or 20x/0.85 NA oil objective lens (OLIMPUS) and a cold CCD camera (CoolSNAP HQ; Roper Scientific) that was controlled using Delta Vision SoftWorx (Applied Precision). The Z-stack images were then deconvoluted.

### Image analysis

To quantify the spindle rotation angle, the centroid of each separated chromosome (10 min after Ana-II onset before the beginning of spindle rotation), the egg, and the spindle midpoint were tracked by image processing using ImageJ. For the chromosome centroid, fluorescence images of the chromosomes (H2B-mRFP1) were smoothed with a median filter (radius = 2) and binarized using a threshold (Yen algorithm). For the egg centroid, the cytoplasmic fluorescence images of Cal-520-dextran or EGFP-α-tubulin or H2B-mRFP1 were smoothed with a median filter (radius = 10), then binarized using a threshold (Otsu algorithm). The centroid coordinates of the binarized images, defined as the x-y coordinates, were calculated using ImageJ. The spindle midpoint was defined as the midpoint between the centroids of the separated chromosome signals. The spindle rotation angle was quantified as the angle between the line connecting the centroids of the separated chromosome signals and the line connecting the egg centroid and spindle midpoint. The coordinates of the chromosome centroids at n min after Ana-II onset were defined as A (*a_n_, b_n_*) for the chromosomes eventually extruded as PB2 and A’ (*c_n_, d_n_*) for the chromosomes remaining in the eggs. The coordinates of the spindle midpoint and egg centroid n min after Ana-II onset were defined as B (*e_n_, f_n_*) and B’ (*g_n_, h_n_*), respectively. The rotation angle is given by

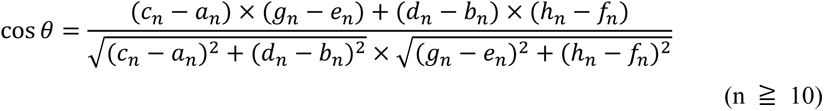

Conversion from radians to angles is given by

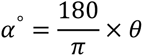

where α° is the rotation angle. The angles were normalized by subtracting all angles from the initial angle to calculate the rotation angle from the initial spindle position. To quantify normalized changes in the rotation angle, the angular velocity (Δt = 1 min) a few minutes before and after the Ca^2+^ transient was divided by the absolute maximum angular velocity for the same period. All Ca^2+^ transients were maintained for 30 min during spindle rotation.

To quantify the spindle position within the egg, the distance (*d*) between the spindle midpoint and egg centroid was calculated. The coordinates of both the spindle midpoint and egg centroid were obtained using the same method as that used to quantify the spindle rotation angle.

Ca^2+^ oscillations were analyzed using the fluorescence signals of Cal-520-dextran. To measure the relative intensity of Cal-520-dextran, its cytoplasmic fluorescence images were smoothed using a median filter (radius = 10) and binarized using a threshold (Otsu algorithm). After defining a region of interest (ROI) using the binarized images of Cal-520-dextran, its mean fluorescence signals within the ROIs were measured. To calculate the relative intensity of cytoplasmic Ca^2+^, Cal-520-dextran signals at each time point were divided by the minimum intensity for 30 min during spindle rotation.

The cortical F-actin intensity in live eggs was measured using EGFP-UtrCH fluorescence signals. A wide range of five pixels of cortical F-actin fluorescence signal was measured as cortical F-actin intensity. To obtain a width of 5 pixels, cytoplasmic fluorescence signals of histone H2B-mRFP1 were smoothed with a median filter (radius = 10) and binarized using a threshold (Otsu algorithm). Second, two binarized images were eroded or dilated so that the difference between the larger and smaller regions was 5 pixels, and ROIs were defined in each image, followed by the measurement of the mean fluorescence intensity of EGFP-UtrCH within the ROIs. Third, the mean fluorescence intensity of the dilated ROI was subtracted from that of the eroded ROI. Three cortical regions were selected for the analysis of cortical F-actin intensity. To calculate the relative intensity of cortical F-actin, the EGFP-UtrCH intensity at each time point was divided by the minimum intensity during 30 min of spindle rotation. For normalization, the relative intensity of cortical F-actin a few minutes before and after the cytoplasmic streaming inversion was divided by the absolute maximum intensity of cortical F-actin for the same period. To validate the cortical myosin II activity in fixed eggs, the fluorescence intensity of pMLC was measured using the plot profile (line width: 20 pixels) in ImageJ. The line for the plot profile was drawn across the center of the egg to avoid the actin cap regions.

To validate the fluorescence intensity of cortical F-actin in the fixed eggs, the average of the two peak values of cortical F-actin intensity, measured using the plot profile (line width: 20 pixels), was calculated after subtracting the background intensity. Ratios of cortical F-actin intensity were obtained by dividing the average intensity of cortical F-actin in eggs with cortical pMLC signals by that in eggs without cortical pMLC signals.

The fluorescence intensities of pMLC and F-actin in the cortical region were analyzed using the fluorescence signals of maximum intensity z-projections (ten z-slices) around the egg equatorial plane.

All heat maps of the normalized values for each image analysis were generated using GraphPad Prism9 software.

### PIV analysis

To analyze cytoplasmic streaming, cytoplasmic particle dynamics were tracked using the PIVlab package^42^. Fluorescent time-lapse images merged with a bright field were used to simultaneously analyze cytoplasmic streaming with spindle rotation and Ca^2+^ oscillations. Because the egg in the time-lapse images moved slightly in the x-y plane, the StackReg plugin written for ImageJ was used to stabilize the position of the eggs before PIV analysis. The non-egg area was masked to exclude vectors outside the egg. We used the following parameters for PIV analysis, similar to those previously described^19,20^: CLAHE window size: 100 pixels, high-pass kernel size: 10 pixels, interrogation area of pass 1: 70 pixels, interrogation area of pass 2: 35 pixels, sub-pixel estimator: Gauss 2x3-point, correlation robustness: standard. The analyzed vector maps were adjusted and smoothed using vector validation and modification functions, respectively. For display purposes, the velocity magnitude of cytoplasmic streaming in the y-axis direction of the x-y plane (the axis perpendicular to the spindle) was colored using the v-component of the display function. The blue color indicates a positive value of streaming toward the spindle, and the yellow color indicates a negative value of streaming opposite the spindle. After obtaining vector maps of consecutive frames, the mean value of the vectors within a specific square area below the spindle was calculated as the velocity and direction of cytoplasmic streaming, as shown in Fig 2**a,d**. To quantify the normalized velocity of cytoplasmic streaming, the velocity of cytoplasmic streaming minutes before and after the Ca^2+^ transient was divided by the absolute maximum velocity during the same period. All Ca^2+^ transients were selected for 30 min during spindle rotation. Heat maps of normalized cytoplasmic streaming were created using the GraphPad Prism9 software.

### Quantifications and statistical analysis

Statistical analyses were performed using R or GraphPad Prism9. First, the normality of the data was checked using the Shapiro–Wilk test. The data shown in Extended Data Fig. 3a were parametric and tested using Welch’s t-test. The data in Fig. **7b,g** are non-parametric and were tested using the Mann–Whitney *U* test. For representative images, experiments were performed at least thrice. All experiments involving eggs were performed at least two or more times, where n denotes the number of eggs.

## Data availability

The data that support the finding of this study are available from the authors upon request.

## Acknowledgements

We thank Tomo Kondo (The University of Tokyo) for advice on setting up a holing pipette, Kazuo Yamagata (Kindai University) for advice on the time-lapse imaging of mouse eggs, Atsuo Ogura (RIKEN BRC) and Junya Ito (Azabu University) for the human-PLCζ constructs, Hiroshi Kimura (Tokyo Institute of Technology) for the mRFP1-tagged histone H2B constructs, Bill Bement (University of Wisconsin-Madison) for the EGFP-UtrCH constructs. The research was supported by the Japan Society for the Promotion of Science KAKENHI (grant numbers 21J11440 to T.T.; 20H03250, 20H05356, 22H04664 and 23H02485 to M.O.).

## Author contributions

T.T. and M.O designed the research. T.T. performed all experiments and data analysis. T.T. and M.O. wrote the manuscript.

## Extended Data Figure legends

**Extended Data Fig.1.**
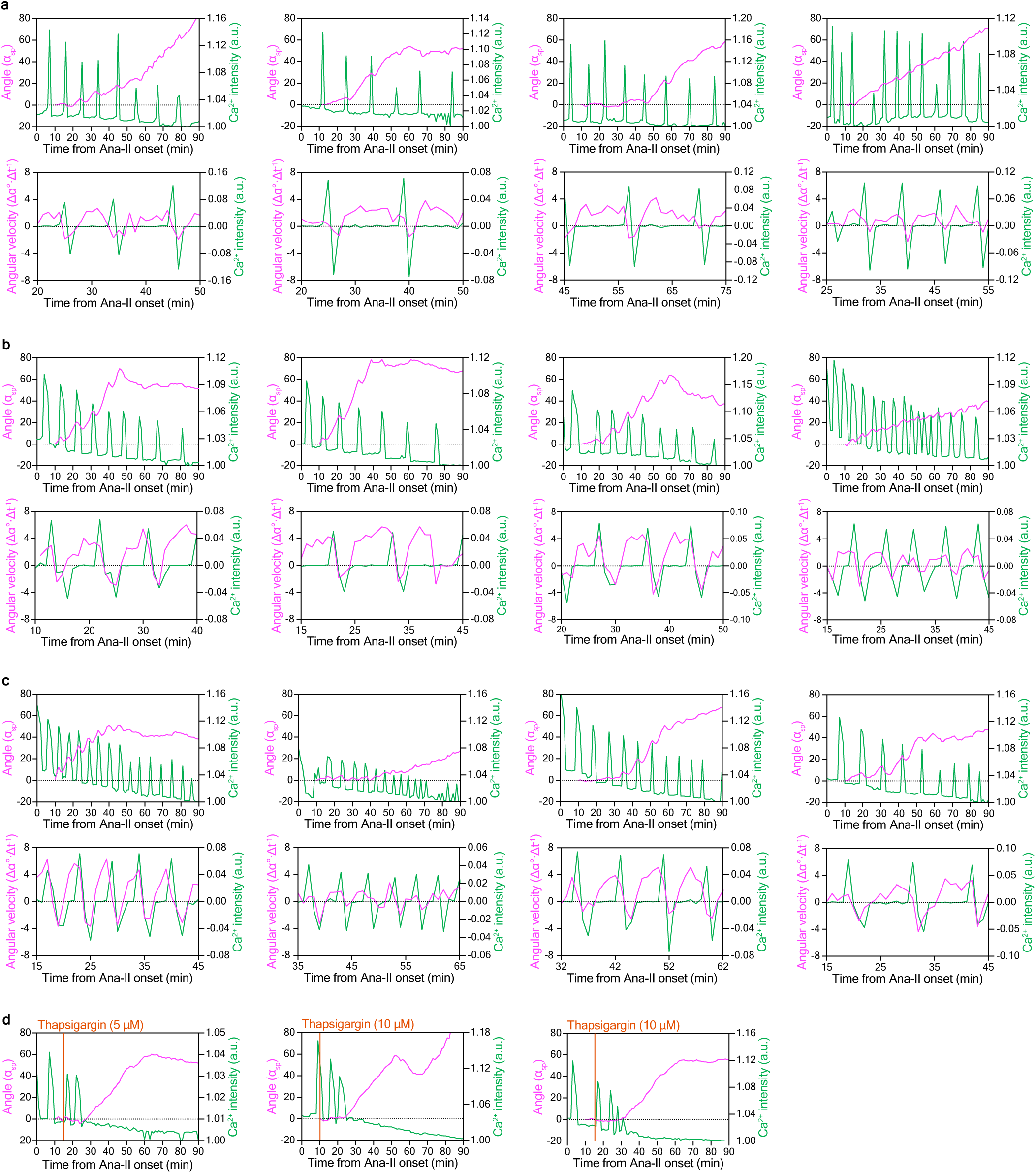
Transient inversions in the direction of spindle rotation is dependent on Ca^2+^ oscillations. **a,b,c,** Graphs showing the spindle rotation angle (magenta) and relative intensity of cytoplasmic Ca^2+^ levels (green) (top panels), and the angular velocity of (Δt = 1 min) of spindle rotation and changes in the relative intensity of cytoplasmic Ca^2+^ for 30 min during spindle rotation (bottom panels) in four representative IVF **a**, or Sr^2+^-activated **b,** or human-PLCζ expressing Sr^2+^-activated **c** eggs. **d,** Graphs showing the spindle rotation angle (magenta) and relative intensity of cytoplasmic Ca^2+^ levels (green) in three representative thapsigargin-treated eggs. Orange bars indicate the time of thapsigargin addition.

**Extended Data Fig.2.**
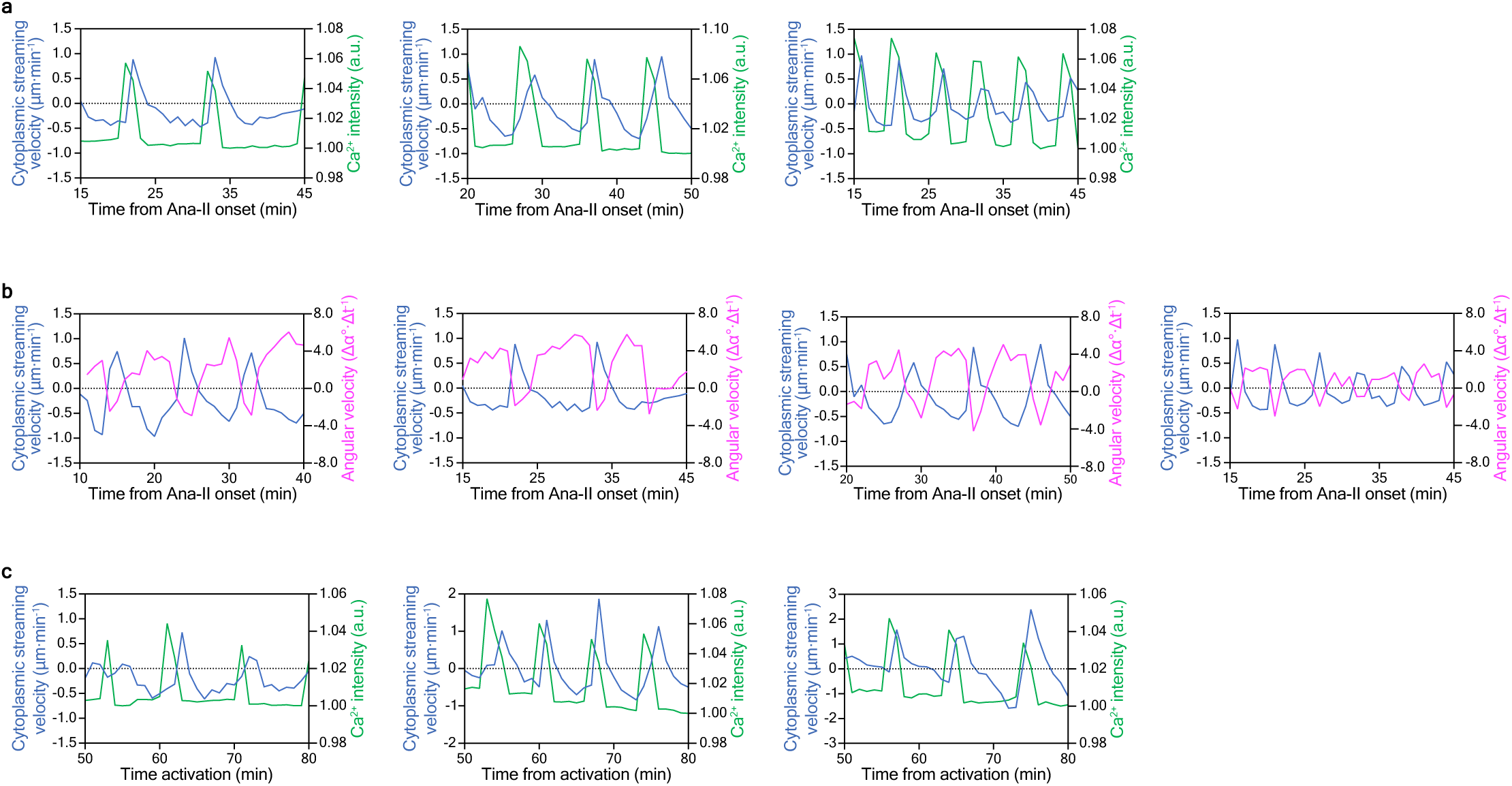
Inversion of cytoplasmic streaming immediately after the Ca^2+^ transient cause a reversal of spindle rotation. **a,c,** Graphs showing the velocity of cytoplasmic streaming (blue) and relative intensity of cytoplasmic Ca^2+^ levels (green) in three representative Sr^2+^-activated eggs **a**, or Sr^2+^-activated eggs treated with 0.08 µg/mL nocodazole **c**. **b,** Graphs showing the velocity of cytoplasmic streaming (blue) and angular velocity of spindle rotation (Δt = 1 min) (magenta) in four representative Sr^2+^-activated eggs.

**Extended Data Fig.3.**
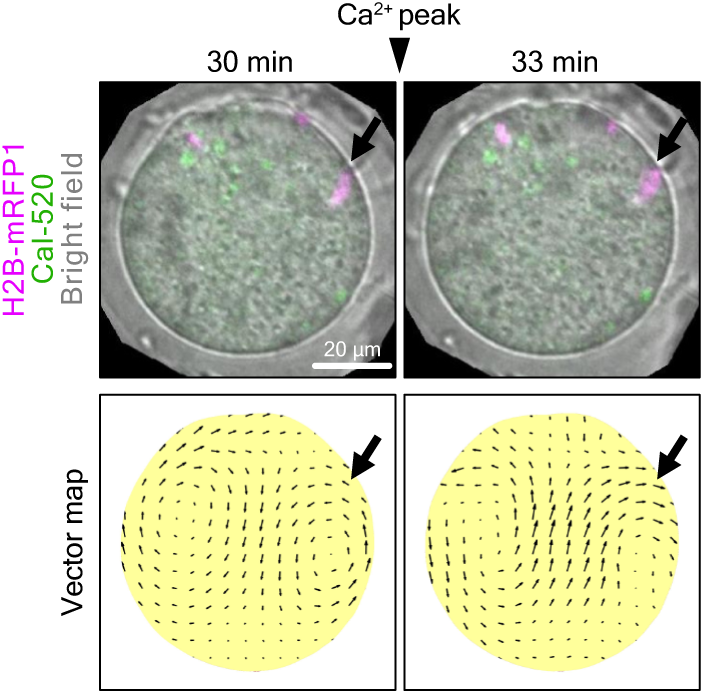
Inversed cytoplasmic streaming is oriented towards both maternal and paternal chromosomes. Representative images showing chromosomes (magenta) and cytoplasmic Ca^2+^ levels (green) in an IVF egg at a time point before and immediately after the Ca^2+^ transient merged with a bright field (top panels). Vector maps indicating the direction and velocity of cytoplasmic streaming analyzed using PIV (bottom panels). Black arrows indicate the positions of sperm chromosomes. Numbers above the top panels indicate the time after the onset of Ana-II.

**Extended Data Fig.4.**
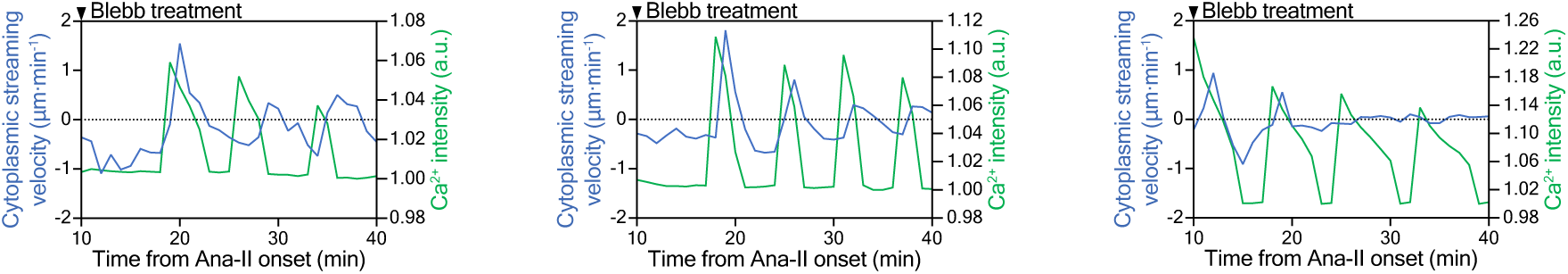
Inhibition of myosin II activity decrease the velocity of cytoplasmic streaming but not compromise Ca^2+^ oscillations. Graphs showing the velocity of cytoplasmic streaming (blue) and relative cytoplasmic Ca^2+^ levels (green) in three representative blebbistatin-treated Ana-II eggs. Black arrowheads indicate the time of blebbistatin addition.

**Extended Data Fig.5.**
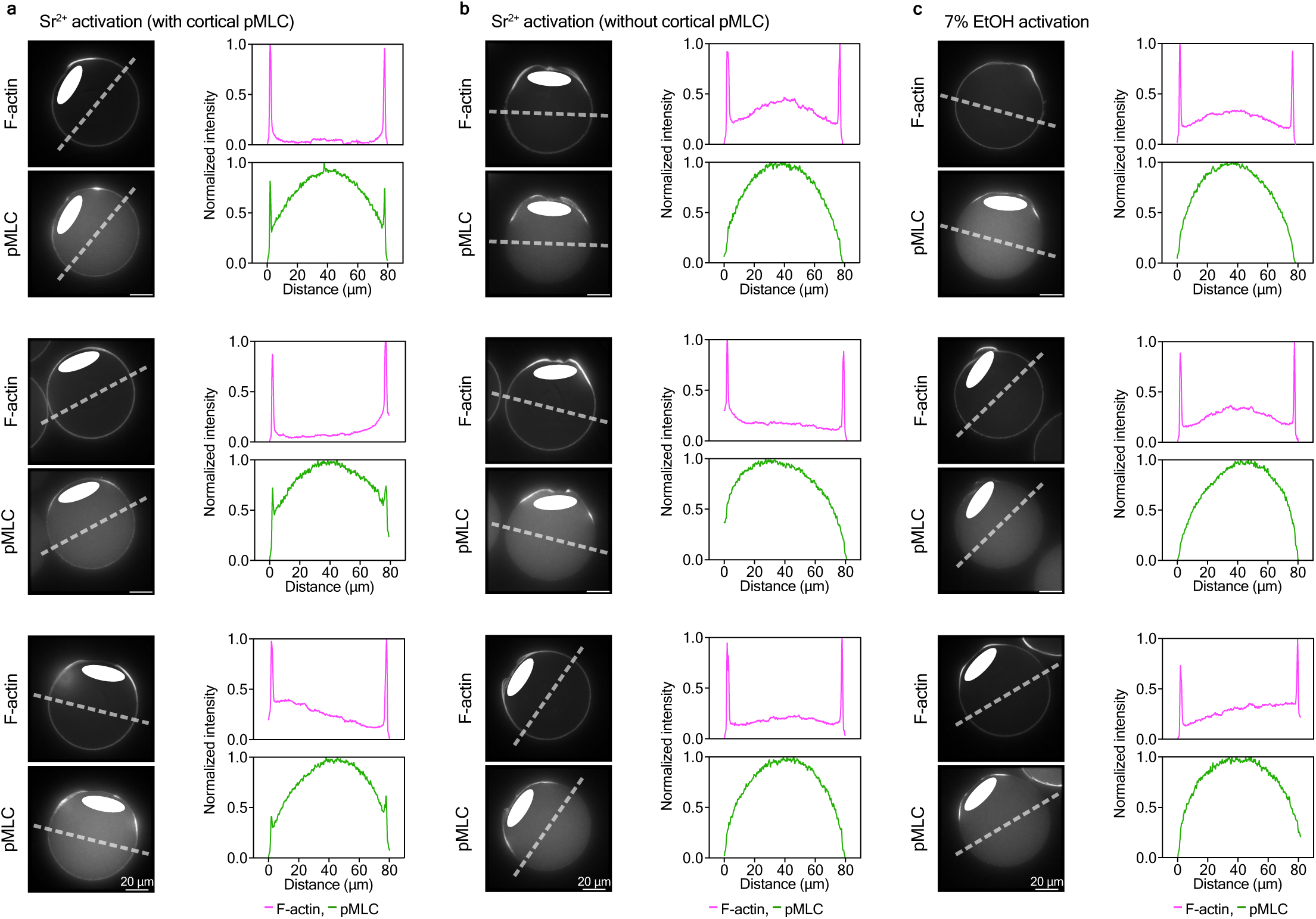
MLC phosphorylation at the cortex in Sr^2+^ activated eggs. **a,b,** Left, three representative fluorescent images of F-actin and pMLC in Sr^2+^ activated eggs with **a** or without **b** cortical pMLC singals. Right, graphs show the fluorescence intensity profiles of F-actin and pMLC along the dashed white lines in the left panels. **c,** Left, three representative fluorescent images of F-actin and pMLC in 7% EtOH activated eggs. Right, graphs show the fluorescence intensity profiles of F-actin and pMLC along the dashed white lines in the left panels. White ellipses indicate the Ana-II spindle.

